# Reconstructing fresh green leaf spectra in the SWIR-2 region (2001–2500 nm) collected in a humid environment by referring to publicly available green leaf spectral databases

**DOI:** 10.1101/2024.01.04.574267

**Authors:** Lino Garda Denaro, Hsin-Ju Li, Jie-Yun Chong, Cho-ying Huang

## Abstract

Leaf spectra (reflectance and transmittance) are key parameters for land surface physical and biogeochemical modeling and are commonly measured using a portable spectroradiometer and an integrating sphere or contact probe with an artificial light source. However, spectral data may be obscured mainly because of water vapor and low signal-to-noise ratios, especially in the shortwave infrared-2 region (SWIR-2, 2001–2500 nm). This erroneous pattern is particularly pronounced in humid conditions, such as in many tropical and subtropical regions, making data unusable in SWIR-2. In this study, we proposed a statistical/mathematical spectral reconstruction approach to retrieve noise-free SWIR-2 fresh green leaf spectra by referring to the available previously published quality-controlled fresh green leaf reflectance and transmittance reference databases. We processed 896 pairs of fresh tea (*Camellia sinensis* var. *sinensis*) leaf reflectance and transmittance data from Alishan in central Taiwan. The spectral data were acquired by a field spectroradiometer with an integrating sphere. We selected a subset (500–1900 nm) of the spectra in the visible, near-infrared, and SWIR-1 regions (VNS-1) that was relatively insensitive to atmospheric conditions. Then, we applied a Gaussian fitting function to smooth the spectral profile. We matched those spectra with publicly available, quality-controlled, and Gaussian fitting function smoothed reference green leaf spectral databases obtained from Italy (LOPEX), Panama (SLZ), and Puerto Rico (G-LiHT) (1694 reflectance and 997 transmittance samples) and selected the one that was most similar (yielding the highest correlation coefficient) to each smoothed Alishan VNS-1 spectrum. We then used multivariable linear regression, linear parameter multiplication, and spectral reversion to reconstruct SWIR-2 spectra based on VNS-1 spectra. To assess the validity of the proposed SWIR-2 reconstruction method, we acquired an independent set of green leaf spectral databases from France (Angers) with SWIR-2 of 2001– 2450 nm. We found that the performance of the SWIR-2 reconstruction approach was satisfactory, with mean (± standard deviation) root-mean-square errors (RMSEs) of 0.0041 ± 0.0019 (reflectance, 3.0% of the mean SWIR-2 of the test data) and 0.0054 ± 0.0027 (transmittance, 2.5%) for each spectrum and RMSEs of 0.0058 ± 0.0027 (reflectance, 4.2%) and 0.0055 ± 0.0043 (transmittance, 2.5%) for each SWIR-2 band. The proposed approach successfully modeled SWIR-2 of the test spectra, which could be further improved with the availability of a more comprehensive set of green leaf reference spectral databases.

## 1. Introduction

Leaf reflectance and transmittance (collectively defined as “leaf spectra” and used interchangeably in this study) are primary biophysical parameters of the terrestrial environment to model the bidirectional reflectance distribution function (BRDF) and corresponding hemispherical integrated expressions, the hemispherical-directional reflectance factor (HDRF) (Strub et al. 2002), of vegetated land surfaces (Li and Strahler 1985, Ni et al. 1999). These spectral parameters can be used to assess the Earth’s energy budget (Liang et al. 2019). Leaf spectra can also be used to indirectly infer plant biophysical and chemical properties or traits from chemometric analysis (Asner and Martin 2009). One of the most common approaches to measuring leaf reflectance and transmittance is to use a portable field spectroradiometer with an integrating sphere (measuring HDRF) or a contact probe (bidirectional reflectance factor [BRF]) (Asner 1998). However, data are susceptible to air humidity inducing random noise, particularly in the water absorption bands at 1351–1434 nm, 1801–1960 nm, and 2451–2500 nm within the optical region (400–2500 nm). Spectra measured in a laboratory environment with an artificial light source can reduce the impacts on the former two regions. However, because of the low energy source (a low signal-to-noise ratio [SNR]), particularly at a longer wavelength (referring to Planck’s blackbody radiation law) (Planck 1991), pronounced errors may occur in the second shortwave infrared region (SWIR-2, 2001–2500 nm) (see examples in Noda et al. 2014, Simic et al. 2014). For fresh leaf tissues, the spectral variations in this region are particularly sensitive to leaf water content (Curran 1989, Huang et al. 2019a); the errors in SWIR-2 make it difficult to assess.

One commonly used approach to suppressing the noise in SWIR-2 is to apply a filter, such as the Savitzky-Golay (Savitzky and Golay 1964) or simply low pass (Wang and Sousa 2009) filters, to smooth erroneous data patterns of leaf spectra (He et al. 2007, Ali et al. 2016). Schmidt and Skidmore (2004) applied wavelets, which have also often been used to smooth leaf spectra (Kokalj et al. 2011, Sexton et al. 2021). However, these approaches may be ineffective if data noise is substantial, particularly for the SWIR-2 bands (see the examples of Noda et al. 2014, Qi et al. 2014, Roelofsen et al. 2014, Potůčková et al. 2016, Mõttus et al. 2017, Hovi et al. 2021).

Returning to the discussion of Planck’s radiation law, the energy of a blackbody is governed by temperature and wavelength. The peak radiant exitance (W m^-2^ nm^-1^) for the sun (5778 K) is at approximately 450 nm in the blue region. The energy drops abruptly with increasing wavelength toward 2500 nm. Therefore, spectra are less error-prone in the visible, near-infrared (NIR) and SWIR-1 (VNS-1, 400–2000 nm) regions because of a greater SNR compared to SWIR-2 bands. Leaf spectra may be influenced by several plant physical and physiological traits (Kokaly et al. 2009, Ustin et al. 2009). Leaf optics can be partitioned into six subregions reflecting the responses of plant physiology and biochemistry: Visible (400–680 nm), red edge (681–740 nm), NIR shoulder (741–900 nm), NIR plateau (901–1300 nm), shortwave-infrared 1 (SWIR-1: 1301–2000 nm) and shortwave-infrared 2 (SWIR-2: 2001–2500 nm) spectral regions (Huang et al. 2019a). The spectra in the visible subregions were dominated by photosynthetic (mainly chlorophyll a and b, carotenoids and xanthophylls) and nonphotosynthetic (anthocyanins) pigment absorption related to photosynthesis (Ustin and Gamon 2010). The derivatives of the reflectance and the wavelength position of the red edge may be related to chlorophyll content and environmental stress, respectively (see the synthesis of Ustin et al. (2009) for details). Spectral signals contributed by solar-induced fluorescence emission peaks at 685 nm and 730 nm (Mohammed et al. 2019) are minor (∼2% of solar radiation) and are not discussed here. Overall, the leaf spectra with NIR regions (the shoulder and plateau, 740–1300 nm) are related to cell structure (e.g., the spatial characteristics of the mesophyll cells and the fraction of air space (Jacquemoud and Ustin 2019)), except for the water absorption “dips” on the plateau centered at 980 nm and 1160 nm caused by the chemical bonds in the biochemical of interest. The spectra of SWIR (1300–2500 nm) are generally inversely proportional to leaf water content, especially at approximately 1400 nm, 1900 nm and 2500 nm (Huang et al. 2019a), with a secondary influence from dry matter in dry leaves (Kokaly et al. 2001, Kokaly et al. 2009).

The “shape” of healthy green vegetation in the optical region possibly can be modeled (e.g., SWIR spectra may be a function of the absorption coefficient of pure liquid water) (Jacquemoud and Baret 1990, Feret et al. 2008, Jacquemoud et al. 2009). According to the previous section, we also understand that the spectral characteristics of green leaves are sensitive to leaf cellular structure and physiology. Therefore, we assume that two healthy green leaf spectra with similar VNS-1 patterns should have similar characteristics of SWIR-2. This assumption may allow us to reconstruct the spectrum with noisy SWIR-2 through a mathematical transformation by referring to the clean spectrum. Following this, we asked whether we could model a green leaf SWIR-2 spectrum collected in a humid environment based on the similarity of its VNS-1 spectral pattern to reference data from quality-controlled spectral databases.

## 2. Methods and Materials

### 2.1. Test green leaf data acquisition

The primary objective of the study was to reconstruct noisy green leaf SWIR-2 reflectance and transmittance data. For simplicity, we acquired green leaf samples (hereafter “test data” or “test spectra”) from one tea tree species, *Camellia sinensis* var. *sinensis* (but with different cultivars) in 1850 ha of the Greater Alishan tea farms in central Taiwan (Fig. 1). The mean (± standard deviation [SD]) elevation of the tea farms is 1214 ± 398 m above sea level (a.s.l.). According to long-term (1901–2019) climate records (Harris et al. 2020), the mean annual precipitation and air temperature are 2000 mm and 19 °C, respectively. The primary tea products are Chinshin oolong tea (green tip oolong variety) and Jhinhsuan. The soil types in this region are spodosols and ultisols.

**Fig. 1.**
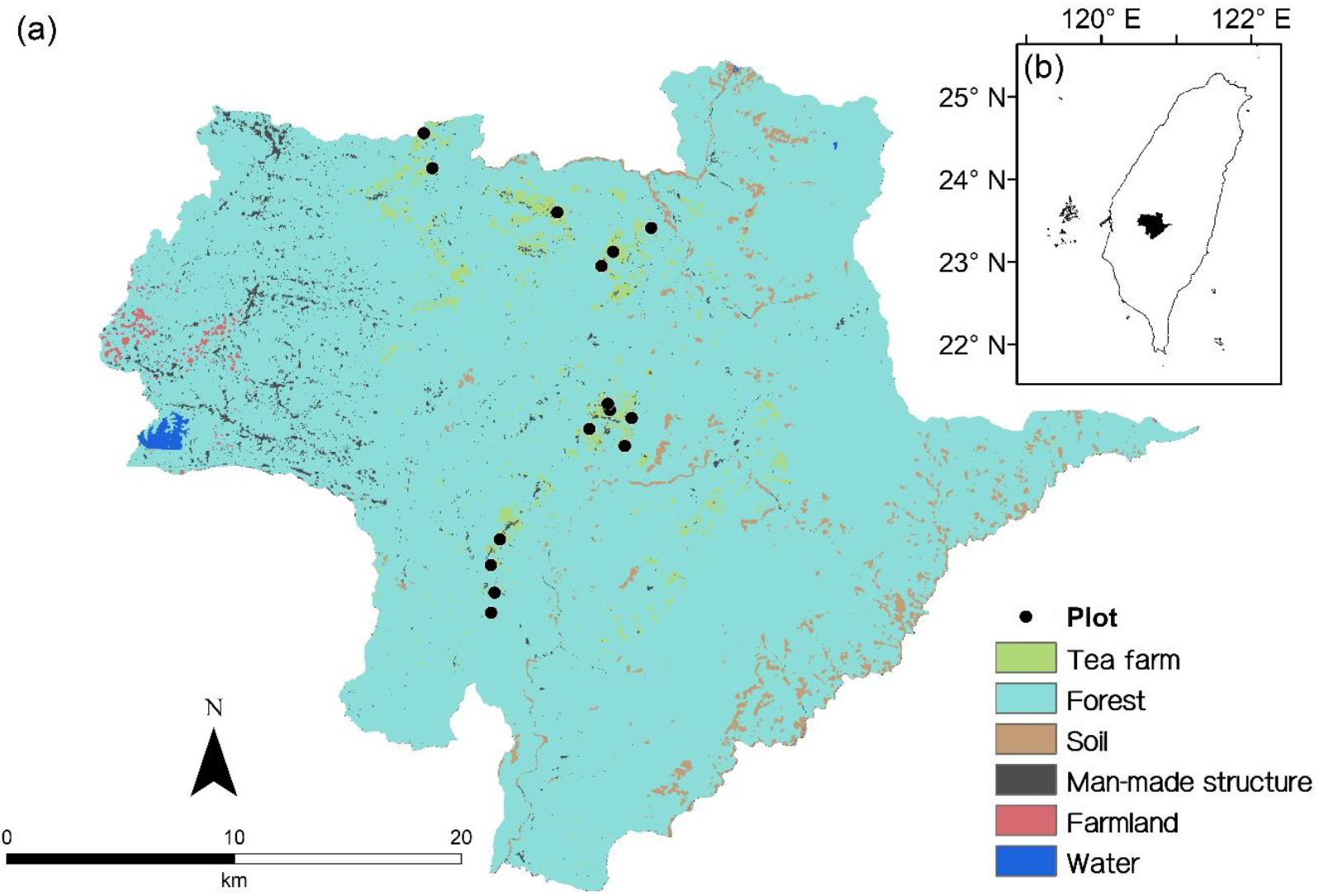
(a) Geographical locations of tea farms for leaf spectral test data acquisition in the Greater Alishan tea country in (b) central Taiwan occupying the Alishan, Fanlu, Meishan and Zhuqi townships. The background of the site map illustrates the primary land use and land cover types of the region.

We collected fresh tea shoots from 15 tea farms across an elevation range of 879–1552 m a.s.l. (mean ± SD = 1225 ± 178 m a.s.l.) (Fig. 1a) from April 7 to June 17 of 2022, covering the spring and summer tea harvesting seasons. We note that tea leaves contain biochemical elements related to tea flavor, such as catechins and caffeine, that are not commonly present in other plants. Since the amounts are much lower than those of other primary biochemical components, such as chlorophylls and leaf water content, to our knowledge, no particular optical spectral bands are sensitive to these tea flavor elements (Yamashita et al. 2021). Therefore, using the spectra of tea leaves should be appropriate for the purpose (to develop a universal methodology) of this study. We randomly collected two airtight polyethylene bags of samples (approximately 60 tea shoot [top-of-canopy] samples per bag), including an apical bud and three leaves, for each farm. Average height (measured by a yard stick) and leaf area index (by LAI-2200C, LI-COR, Inc., Lincoln, Nebraska, USA) were 89.3 cm and 8.7 m^2^ m^-2^, respectively (unpublished data). These samples were stored in a 0 ºC cooler until the spectra were measured, and the time between the sampling and the measurement was less than six hours. We randomly selected 30 leaves for each bag and only measured the spectra of the largest leaf (the third leaf) for each shoot because of the sizes of the sample ports of the integrating sphere (diameters = 13 and 15 mm, for the details of the instrument, see 2.2. Leaf spectral measurement and calculation).

### 2.2. Leaf spectral measurement and calculation

We used a portable spectroradiometer (FieldSpec 3, Analytical Spectral Devices, Inc., Boulder, Colorado, USA) for the tea leaf spectral measurement (test data). The spectral resolutions (full-width-half-maximum) were 3 nm (350–1000 nm) and 10 nm (1000–2500 nm) with sampling intervals of 1.4 nm and 2 nm, respectively. Ten readings were averaged for each spectral measurement to keep leaf samples as fresh as possible for further biochemical analysis. To measure tea leaf reflectance (Rs) and transmittance (Ts) using a nonpolarized measurement, we combined the spectroradiometer and a single integrating sphere (ASD RTS-3ZC, for specifications, see Hovi et al. (2018)) and derived tea leaf spectra following the procedures (eqs. 1 and 2) recommended by the manufacturer to avoid substitution errors (Hovi et al. 2018):

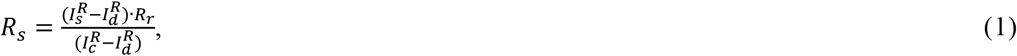

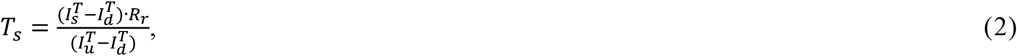

where Is, Ic, Iu and Id are the spectroradiometer readings measured from the tea leaf sample, calibrated and uncalibrated white references, and stray light measurement using a light trap), respectively. Superscripts R and T refer to reflectance and transmittance measurements, and Rr is the reflectance factor of the calibrated white reference. We note that technically, we measured directional reflectance and transmittance factors through the procedure above (Greiner et al. 2007), and the terms leaf reflectance and transmittance were used here for expediency.

### 2.3. Green leaf spectral databases

The SWIR-2 spectral reconstruction approach assumes that SWIR-2 can be modeled based on the data patterns of VNS-1; the possible range and variation in each VNS-1 band must be included. Therefore, we acquired previously published, publicly available and quality-controlled (for example, with detailed documentation) fresh green leaf reflectance and transmittance databases (hereafter “reference data” or “reference spectra”) that covered the entire optical spectral region. The green leaf spectra of conifer species were not included in this study because of considerably different measurement strategies (Hovi et al. 2017, Hovi et al. 2020). Based on these data selection criteria, three reference databases were selected for this study: (i) The Leaf Optical Properties EXperiment (LOPEX) 93 conducted by Istituto Superiore per la Protezione e la Ricerca Ambientale, Italy (Hosgood et al. 1994), (ii) the 2020 San Lorenzo (SLZ) field campaign conducted in the tropical forests of Panama by Brookhaven National Laboratory (Lamour et al. 2021), and (iii) the NGEE (Next Generation Ecological Experiments) Tropics leaf spectral collection in the tropical forests of Puerto Rico (namely, G-LiHT) (Serbin et al. 2019). We visually assessed the data and excluded nongreen vegetation spectra by referring to Asner et al. (2009) and Jacquemoud and Ustin (2019) (Fig. 2, also see Table 1 and Fig. 3 for spectral sample sizes for each green leaf database). Leaf transmittance was measured by single or double integrating spheres, and reflectance was assessed also using those or a contact probe (not specified in metadata) with a 1 nm resampled spectral interval covered the entire optical region from 400 nm to 2500 nm. We note that some minor noises were observed in those spectra; we applied a Gaussian function (Zhao et al. 2021) with a 9 nm window for 40 (to preserve some spectral details) and 100 (to derive overall trends) times to moderately smooth the data for later correlation analysis (Fig. 3).

**Table 1.** Numbers of green leaf spectra (reflectance and transmittance) for the databases to develop (LOPEX, SLZ and G-LiHT) and validate (Angers) the shortwave infrared-2 (SWIR-2, 2001– 2500 nm) reconstruction method. The numbers shown here are before outlier removal. Abbreviations and acronyms of the databases can be found in the main text.

| Database (location) | Reflectance | Transmittance | Spectral range | Reference |
| --- | --- | --- | --- | --- |
| LOPEX (Italy) | 309 | 313 | 400–2500 nm | Hosgood et al. (1994) |
| SLZ (Panama) | 1029 | 461 |  | Lamour et al. (2021) |
| G-LiHT (Puerto Rico) | 356 | 223 |  | Serbin et al. (2019) |
| Alishan (Taiwan) | 896 | 896 |  | This study |
| Angers (France) | 276 | 276 | 400–2450 nm | Feret et al. (2008) |

**Fig. 2.**
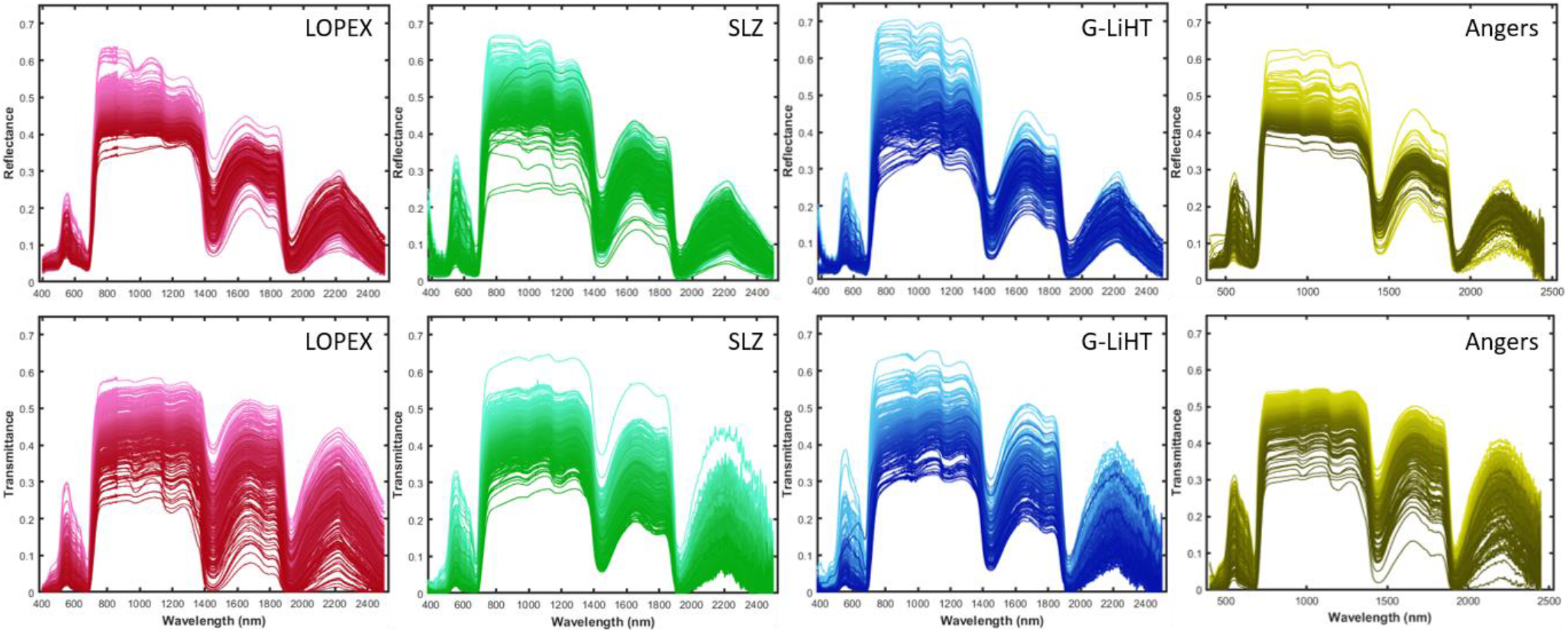
Publicly available green leaf reflectance (upper row) and transmittance (lower row) databases. Subfigures from the same spectral database are illustrated with the same color shades, and there is no corresponding relationship of colors for reflectance and transmittance. The details of the databases can be found in Table 1.

**Fig. 3.**
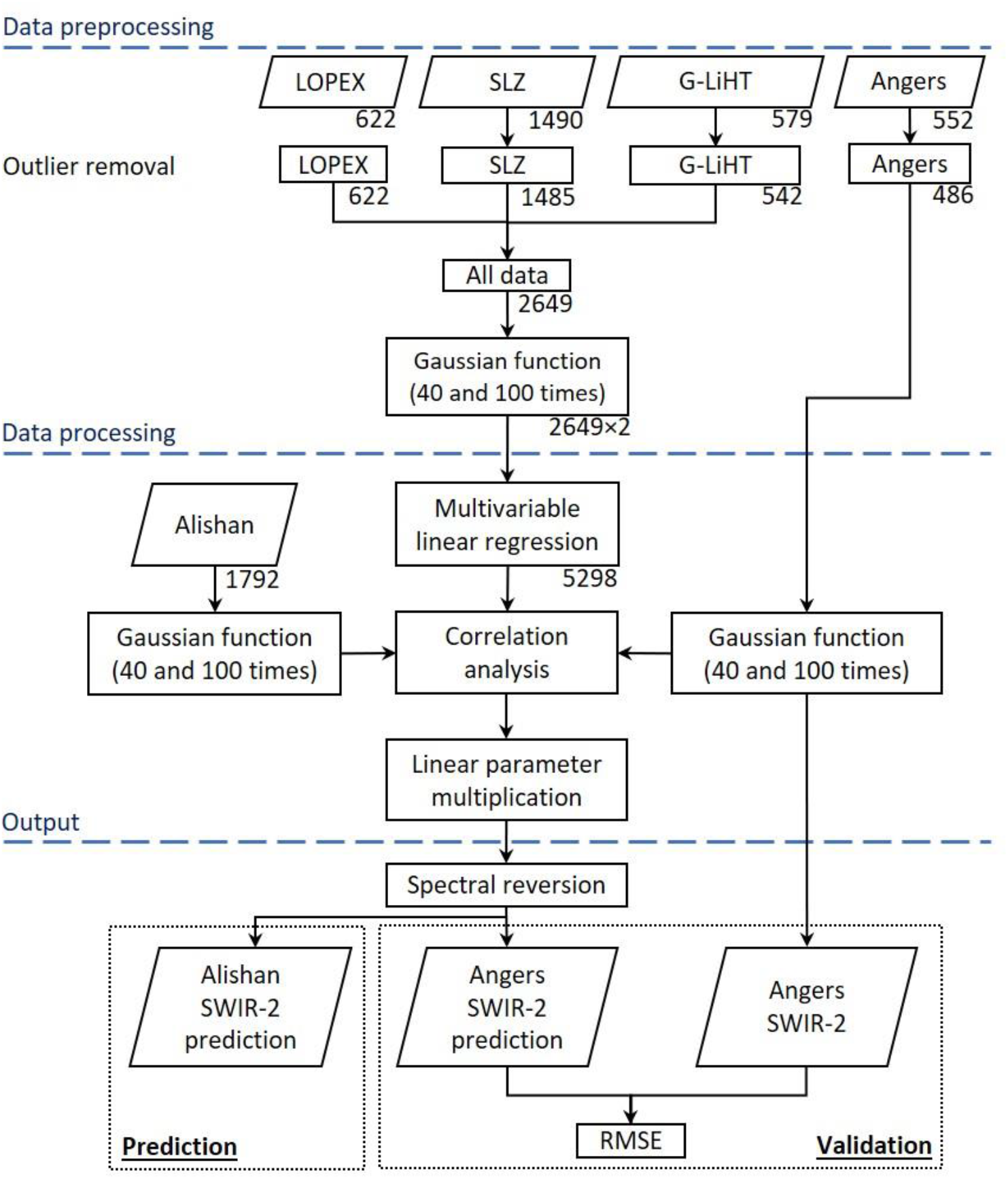
Workflow of the green leaf shortwave infrared-2 (SWIR-2, 2001–2500 nm) spectral reconstruction approach. Parallelograms represent original and final spectral data; rectangles are intermediate products, and numbers outside those boxes indicate sample sizes (reflectance + transmittance, see Table 1 for details).

### 2.4. Reconstruction of SWIR-2 spectra

We employed a series of mathematical and statistical processes to reconstruct noise-prone SWIR-2 spectra. First, for each spectrum in the reference spectral databases (Table 1), we selected the spectral range without apparent noise within VNS-1 to predict its corresponding SWIR-2 using multivariable linear regression. The next step involved preprocessing SWIR-2 spectra of the test data (tea leaf reflectance and transmittance data collected in Alishan) by removing apparent outliers and filling data gaps. We then selected a subset of the spectra (VNS-1) and correlated each VNS-1 spectrum from the test data with all VNS-1 spectra in the reference databases to determine the one with the highest correlation coefficient. After identifying the most correlated reference spectrum, we applied a linear transformation to the test VNS-1 data to model the corresponding SWIR-2 spectra. Finally, we reversed the transformation to finalize the reconstruction. The step-by-step workflow is illustrated in Fig. 3, and further details of these processes are provided below:

The data patterns are relatively erroneous in the ultraviolet-A (350–400 nm) and near the SWIR-2 (1901–2000 nm) regions. Therefore, we only selected 500–1900 nm of VNS-1 (less noise-prone due to higher SNR), to model SWIR-2. The VNS-1 and SWIR-2 wavelengths are written as 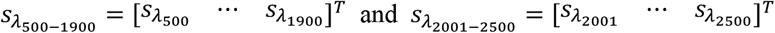, respectively. For this study, we made the assumption that the spectral ranges of the databases (1694 reflectance and 997 transmittance samples) could capture the majority of data variation in green vegetation spectra. We applied a multivariable linear regression to estimate the spectral values at each SWIR-2 wavelength using VNS-1 of 500–1900 nm:

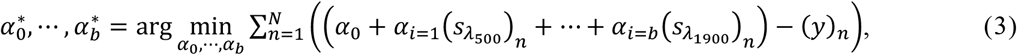

where the 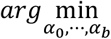 denotes the argument looking for the minimum errors and the optimal parameters denoted as 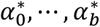, *n* is the index of the number of observation *N*, and *y* is the original SWIR-2 at every band 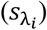; *i* indicates a SWIR-2 band (b) of 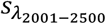. Thus, from the optimization above, the predicted SWIR-2 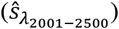 is derived from the linear multiplication of the optimized parameters 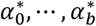 as:

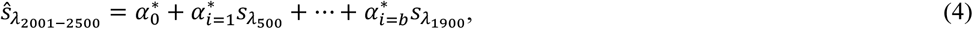

where *i* indicates a spectral number *b* of *VNS*-1 and 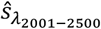 are the predicted of SWIR-2 values 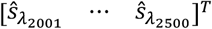. Finally, we smoothed the modeled SWIR-2 spectra with a Gaussian function:

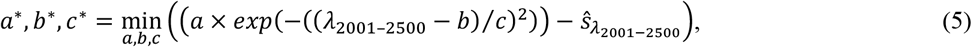

where the 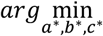 denotes the argument looking for the minimum errors and the optimal parameters denoted as *a*^*\**^, *b*^*\**^, *c*^*\**^, and *λ*_2001−2500_ is the wavelength. Thus, from the optimization above, the predicted SWIR-2 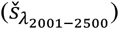 can be derived from the Gaussian function using the optimized parameters *a*^*\**^, *b*^*\**^, *c*^*\**^ (eq. 6):

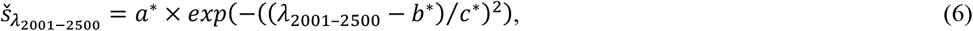

and minimum residual is 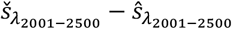. The next step was to preprocess the test data. We first removed apparent outlier data points with values > 1 or < 0 and labeled those data with “null” values. We then filled the data using the mean of spectrally corresponding unlabeled data from all available test data 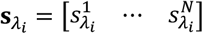, where *N* represents the number of available observations for a particular optical band of 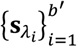; where *i* is the index of spectral band number *b*^*′*^ (500–1900 nm). The gap-filling process for 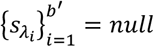 can be formulated as:

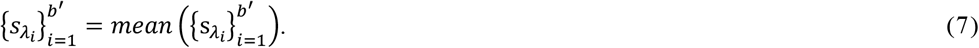

The next step was to moderately smooth the gap-filled test spectral data using a Gaussian fitting function, again with 40 and 100 iterations. We then subset the spectral region of 500–1900 nm 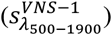 for further analysis.

In the next step, we correlated each test 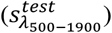 and all reference 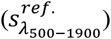 reflectance and transmittance data (smoothed by 40 and 100 times using a Gaussian fitting function) separately to find the one in the databases yielding the highest Pearson’s correlation coefficient.

We then linearly transformed the test data (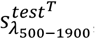, dependent variables) using 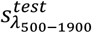 (independent variables):

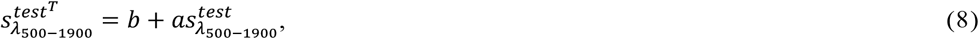

With the coefficients of eq. 3 (a_0_ … a_b_), we could model SWIR-2 of the test data. The modeled SWIR-2 was also moderately smoothed using a Gaussian fitting function (eq. 5). We shifted the modeled SWIR-2 to the corresponding test spectral data by adding or subtracting a constant 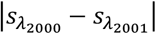 to eliminate the discrepancy. Finally, we reversed the transformed spectra (with shifting 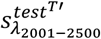) by referring to eq. 8:

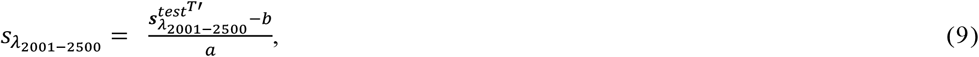

### 2.5. Performance assessment

We used an independent set of green leaf reflectance and transmittance data (Angers) acquired in Angers, France (Feret et al. 2008) from OPTICLEAF (http://opticleaf.ipgp.fr/) to assess the performance of the SWIR-2 reconstruction (using 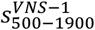 to estimate 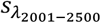). We note that the database was not selected for the reference ones because the spectral data did not cover the spectral range outside 2450 nm, but it should be sufficient for validation. The original dataset contained 276 pairs of reflectance/transmittance data (Table 1). We excluded apparent nongreen vegetation spectra by referring to Asner et al. (2009) and Jacquemoud and Ustin (2019) (Fig. 3). Finally, we derived the root-mean-square error (RMSE) for each spectrum (eq. 9) and each SWIR-2 band of 2001–2450 nm (eq. 10).

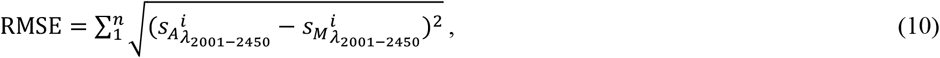

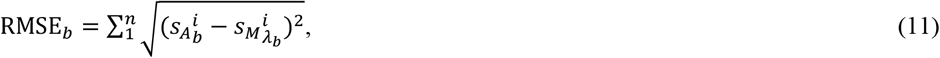

where *s*_*A*_ and *s*_*M*_ are validation (Angers, 243 pairs [a total of 486 samples] for reflectance and transmittance data) and modeled spectra, respectively, and *λ*^*′*^ is each individual SWIR-2 band from 2001 to 2450 nm.

## 3. Results

### 3.1. Test green leaf spectra and the reconstruction of SWIR-2 spectra

In this study, we collected 896 pairs (1792 samples) of green tea leaf reflectance and transmittance from the Greater Alishan tea farms of central Taiwan using a field spectroradiometer and a single integrating sphere. As expected, the spectra were quite noisy based on visual assessment, particularly in the SWIR-2 region (Figs. 4; 7a and 7d). Therefore, we performed the SWIR-2 reconstruction approach. To do so, we needed to preprocess the reference and test data by applying a Gaussian fitting function with 40 and 100 iterations (see Figs. 5 and 6 for illustrations). The data patterns of SWIR-2 of reflectance (Fig. 7b) and transmittance (Fig. 7e) were smooth after reconstruction based on visual assessment. In addition, the mean reflectance and transmittance shifts between the original 2000 nm and modeled 2001 nm were negligible (absolute values < 0.01). There were positive relationships (*p* < 0.001) between the range of absolute (the absolute difference of the original [Fig. 7a and 7d] and modeled [Fig. 7b and 7e] SWIR-2 reflectance and transmittance) errors and wavelengths. In addition, the original spectral values tended to be higher than the reconstructed ones from 2350 nm onward (Fig. 7c and 7f).

**Fig. 4.**
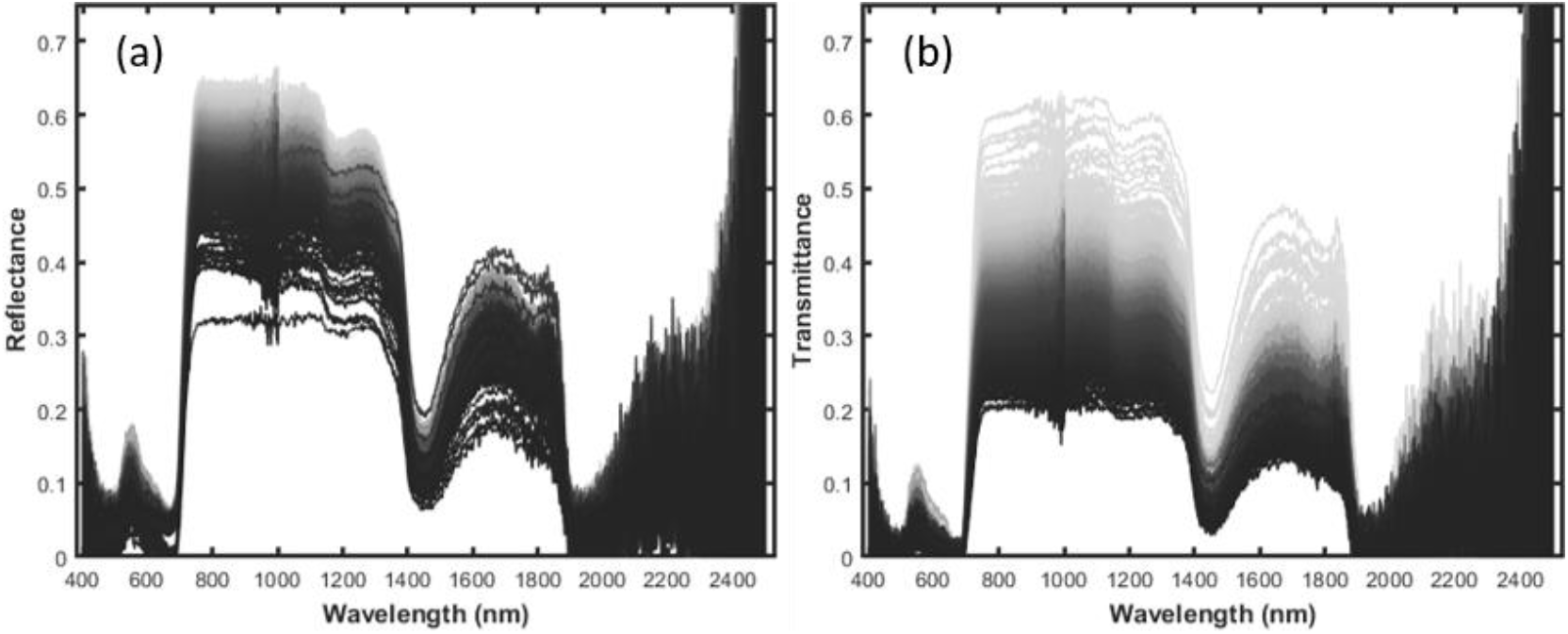
Green leaf (a) reflectance and (b) transmittance data acquired in the tea farms of Alishan in central Taiwan, and there is no corresponding relationship of colors for reflectance and transmittance. The details of the spectral databases can be found in Table 1.

**Fig. 5.**
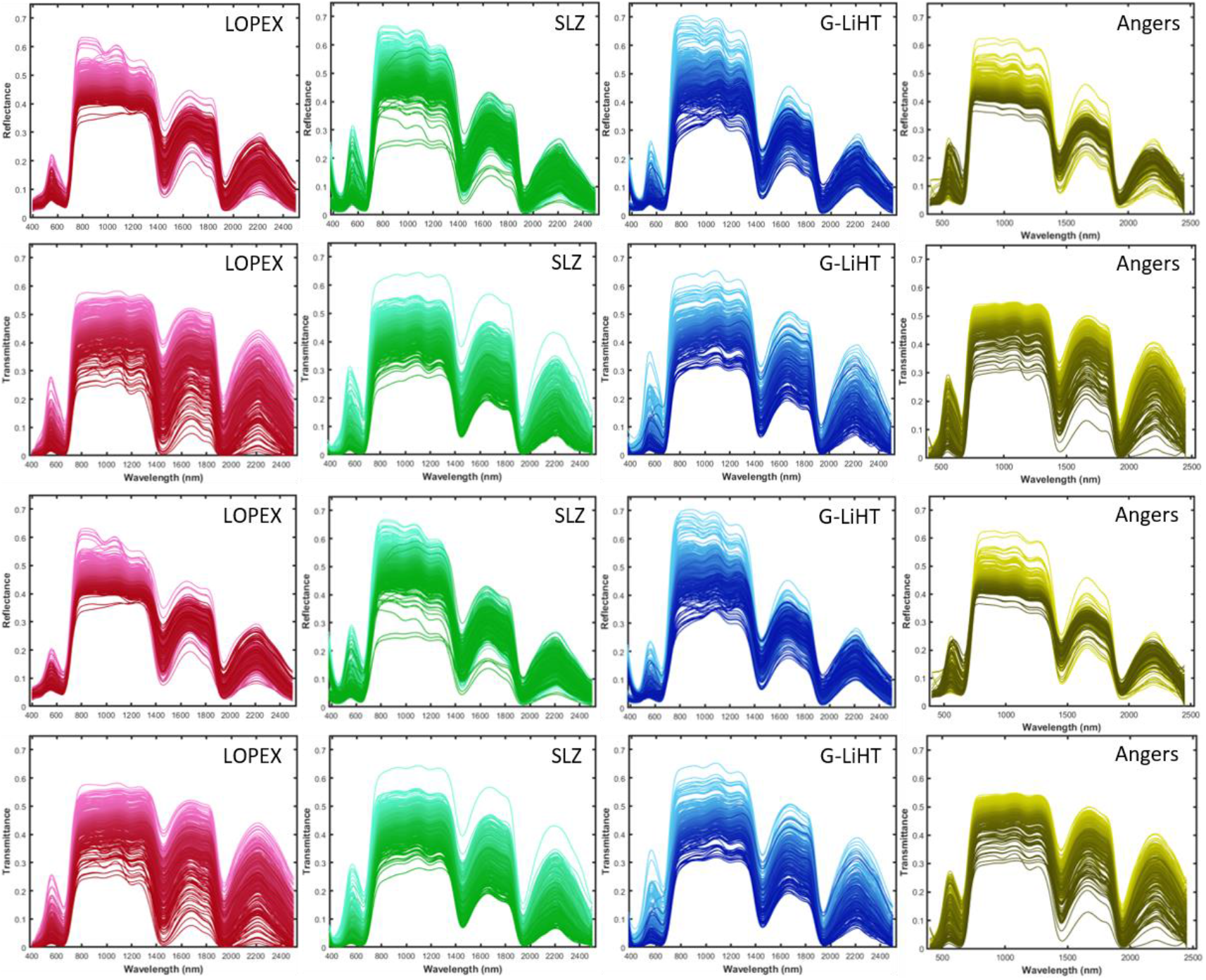
Publicly available green leaf reflectance and transmittance databases smoothed 40 (upper two rows) and 100 (lower two rows) times. Subfigures from the same spectral database are illustrated with the same color shades. The details of the spectral databases can be found in Table 1.

**Fig. 6.**
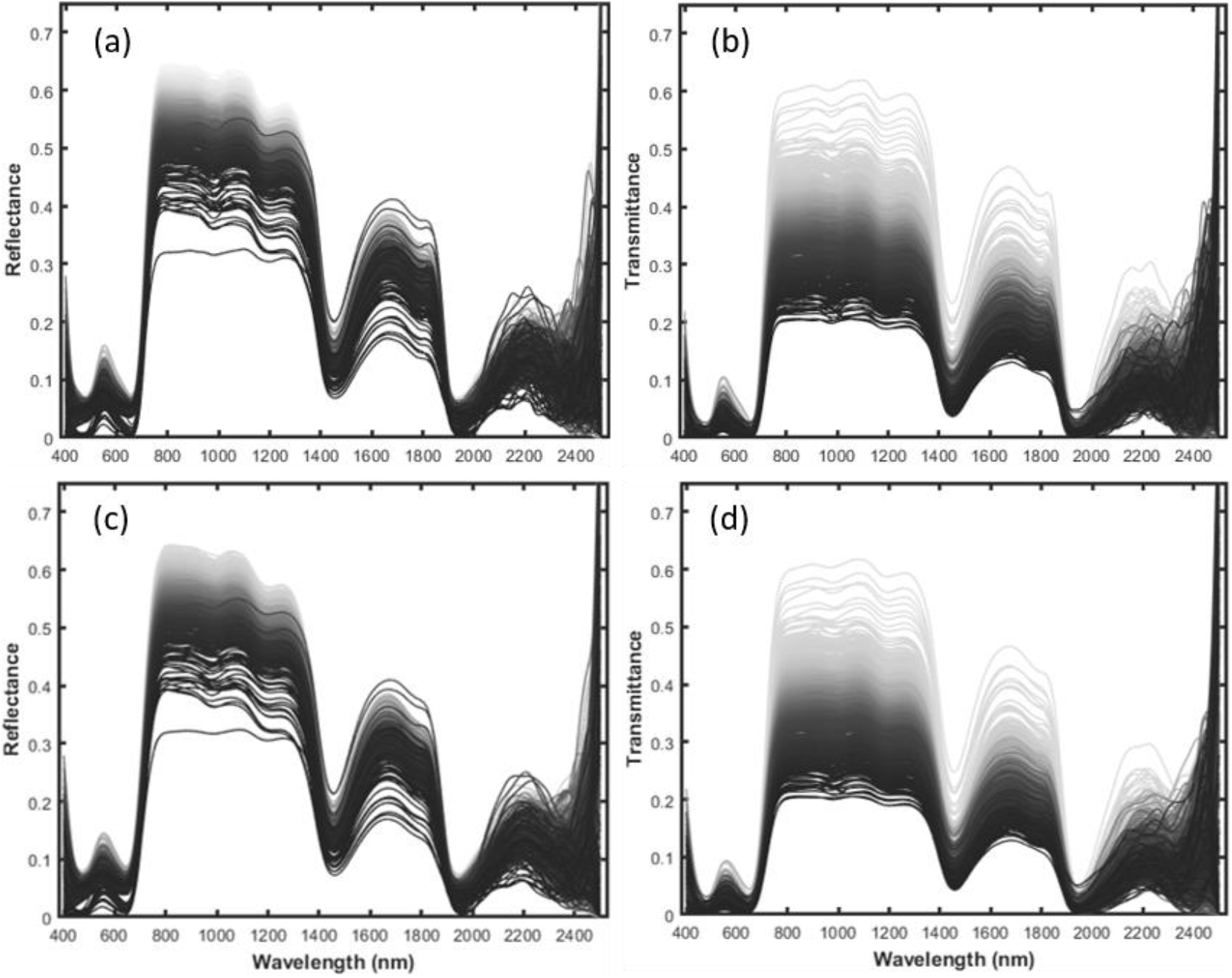
Green leaf spectral data acquired from the tea farms of Alishan in central Taiwan and smoothed 40 (a and b) and 100 (c and d) times. The sample sizes can be found in Table 1.

**Fig. 7.**
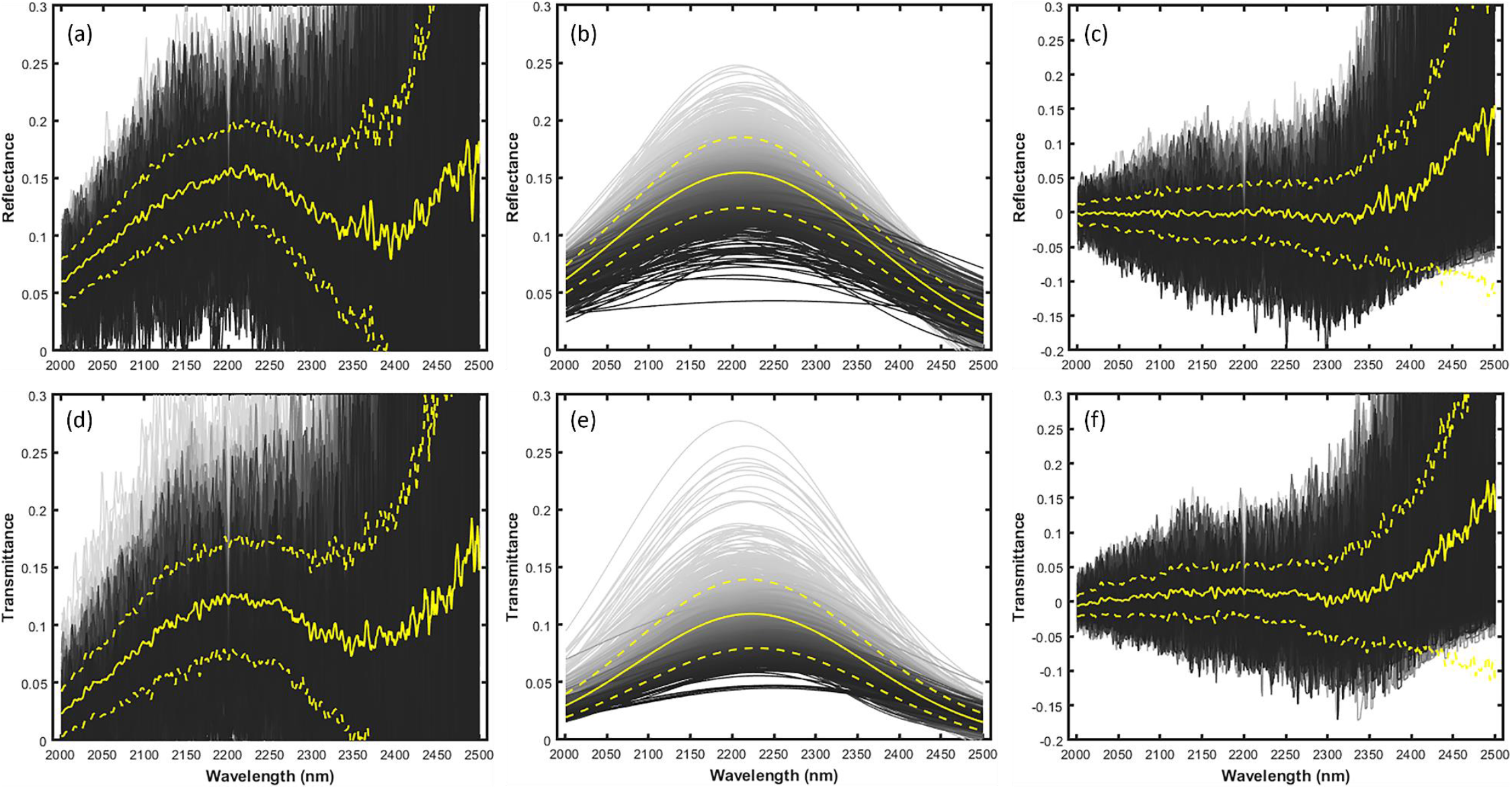
Original (a and d) and reconstructed (b and e) SWIR-2 (2001–2500 nm) Alishan reflectance (upper row) and transmittance (lower row) data and the difference (c and f) between them (c = a - b; f = d - e). The solid and broken yellow lines depict the mean (± standard deviation) of each SWIR-2 spectral band value.

### 3.2. Performance assessment

We used another independent set of green vegetation databases (Angers, Table 1 and Fig. 3) to verify the SWIR-2 reconstruction approach. We found that the performance was satisfactory: for each spectrum (eq. 9), the mean (± standard deviation) RMSE was 0.0041 ± 0.0019 (reflectance, 3.0% of the mean SWIR-2 reflectance of Angers) (Fig. 8a) and 0.0054 ± 0.0027 (transmittance, 2.5% of the mean SWIR-2 transmittance of Angers) (Fig. 8c); for each SWIR-2 band (eq. 10), 0.0058 ± 0.0027 (reflectance, 4.2%) (Fig. 8b) and 0.0055 ± 0.0043 (transmittance, 2.5%) (Fig. 8d). The errors were relatively pronounced at longer bands (e.g., ≥ 2400 nm).

**Fig. 8.**
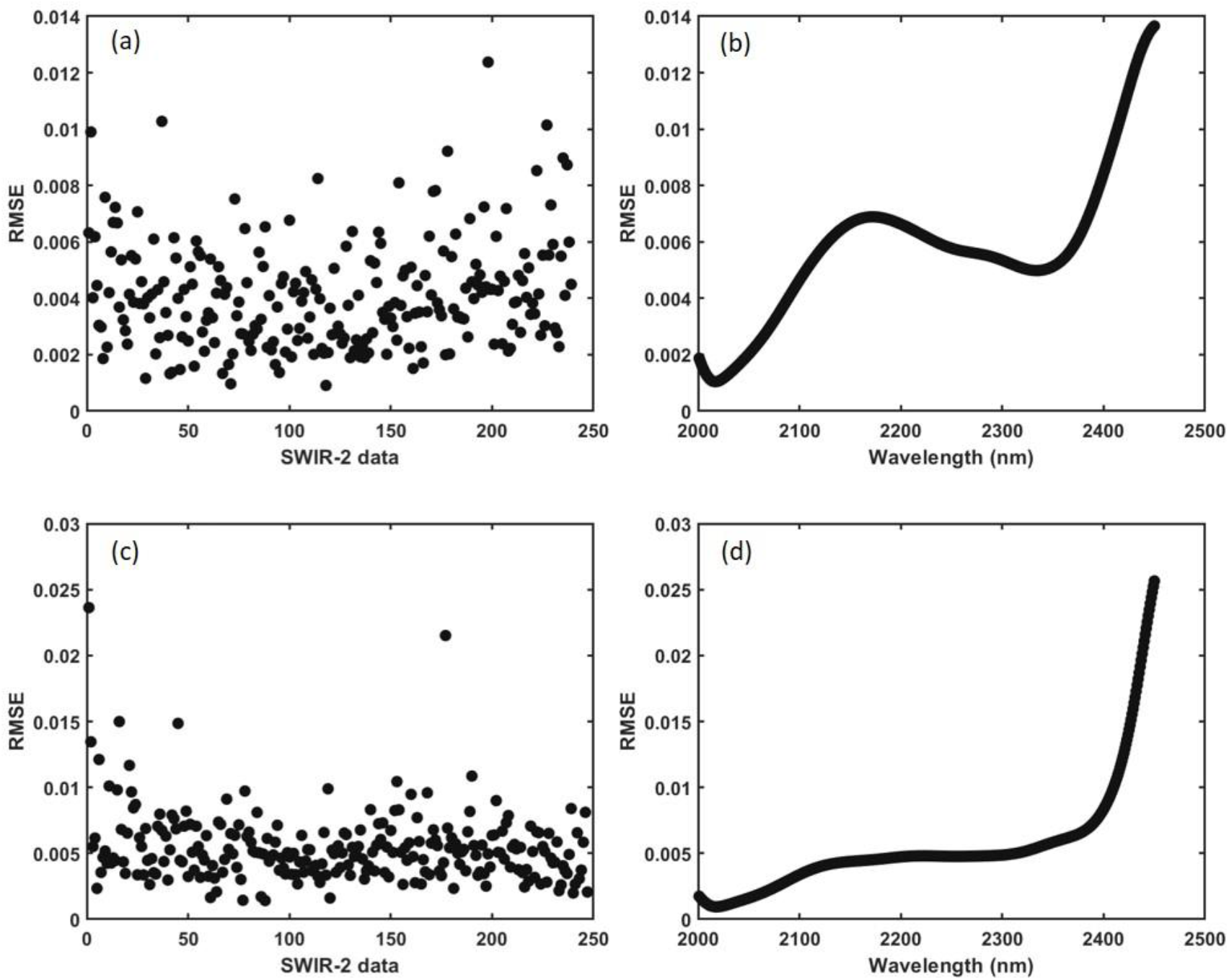
Performance assessment of the SWIR-2 reflectance (upper row) and transmittance (lower row) reconstruction using another independent spectral database (Angers, see Table 1 for details). The validation metrics include the root-mean-square error (RMSE) for each (a) reflectance and (c) transmittance SWIR-2 spectrum and those (RMSEb) for each SWIR-2 (b) reflectance and (d) transmittance band.

## 4. Discussion

### 4.1. Reconstruction of SWIR-2

This study proposed a statistical-based, empirical approach (Fig. 3) to reconstructing the water vapor-affected noisy SWIR-2 (2001–2500 nm) region of green leaf reflectance and transmittance data measured in a humid environment. Overall, the performance was satisfactory (Fig. 8) using an independent set of green leaf reflectance and transmittance databases (Angers) for the assessment. This result may justify the use of the Gaussian function smoothed and less noise-prone 500–1900 nm of VNS-1 of reference and test spectral data (Figs. 5 and 6) to reconstruct those spectra of SWIR-2.

This approach will generate the overall trend of SWIR-2 in green leaves (Fig. 7b and 7e), which is related to leaf water content (Huang et al. 2019a). However, it may not be suitable for modeling the SWIR-2 region of dry leaf samples. Two narrow absorption bands of the reflective characteristic centered at 2054 nm and 2172 nm are sensitive to nitrogen concentration (Kokaly et al. 2001, Kokaly et al. 2009); these subtle features may not be observed after the reconstruction. However, these two narrow absorption bands can be modeled by applying the same approach (using the reflectance of 2054 nm and 2172 nm as the dependent variables) as the secondary process to refine the details of the SWIR-2 reflectance. However, obtaining suitable green leaf spectral databases could be challenging since the SWIR-2 spectral region with minor noise (see examples in Fig. 2) may not be suitable for this specific task.

### 4.2. Spectral noise pattern in the SWIR-2 region

The data noise, defined as the difference between the original and reconstructed spectral values, of the SWIR-2 region of empirical test data acquired in the Greater Alishan tea country in central Taiwan was amplified with increasing wavelength (Fig. 7a and 7d). In addition, the patterns may not be random and move upward rapidly after 2350 nm for reflectance and transmittance (Fig. 7c and 7f), possibly due to the low SNR. This behavior would limit the feasibility of commonly used data smoothing methods, such as spectral filtering including the Savitzky-Golay (Savitzky and Golay 1964), a low pass filter (Wang and Sousa 2009), or a wavelets approaches (Schmidt and Skidmore 2004). These methods generate biased data toward the end of SWIR-2 (see Fig. 6 for an example). The modeled green leaf SWIR-2 spectra by the proposed reconstruction method are constrained by *a priori* knowledge from previously acquired reference databases (Fig. 2) (Hosgood et al. 1994, Serbin et al. 2019, Lamour et al. 2021), which minimizes this erroneous trend.

### 4.3. Future improvement and directions

The performance of SWIR-2 spectral reconstruction was highly dependent on the comprehensiveness of the reference databases. As much spectral variation in VNS-1 vs. SWIR-2 as possible was included so that the SWIR-2 spectral reconstruction approach could interpolate but not extrapolate the test data. In this study, we compiled three green leaf spectral databases of trees and crops from Italy, Panama, and Puerto Rico (Table 1 and Figs. 2 and 3). Therefore, the proposed method could be further improved with the availability of more high-quality spectral data. However, the challenge could remain to measure the leaf spectral reflectance and transmittance of colored fresh plant tissues (see Huang et al. 2019b for example) with relatively abundant anthocyanins, betalains, carotenoids and/or xanthophylls without including those in the reference databases. The SWIR-2 reconstruction approach should also work for stressed green leaf spectra since they show two narrow water absorption regions within the NIR plateau centered at 980 nm and 1160 nm with higher reflectance/transmittance (or shallower dips) (Asner et al. 2016, Huang et al. 2019a). In summary, we conjecture that the SWIR-2 reconstruction approach should be an effective tool for different green leaf conditions, but further investigation is indeed necessary. The proposed approach may be used to improve the quality of the relatively weak SWIR-2 signals of vegetated land surfaces acquired by hyperspectral satellite imagery such as the Environmental Mapping and Analysis Program (EnMAP) (Okujeni et al. 2015). Finally, several airborne narrow-band multispectral (Huang et al. 2019b) or hyperspectral (Asner et al. 2007) sensors restrict the spectral ranges of optical remotely sensed data collection to the visible and NIR regions because of the hardware (a silicon array for visible and NIR regions and an InGaAs photodiode for SWIRs-1 and -2 regions) and perhaps budget limitations. With a valid plant physiological theory to support the direct and indirect relationships (Huang et al. 2019a, Jacquemoud and Ustin 2019), the green canopy reflectance of SWIR-1 and/or SWIR-2 spectral regions possibly could be modeled based on the variation in those spectra in the visible and NIR regions.

## 5. Conclusions

Obtaining high-quality leaf reflectance and transmittance is challenging in humid conditions, particularly in the noise-prone SWIR-2 region, mainly due to the relatively low signal and the effect of water vapor. In this study, we proposed a statistics-based leaf SWIR-2 spectral reconstruction approach. Since the reflectance and transmittance of SWIR-2 of fresh green leaves are directly related to leaf content and possibly plant health, we assumed that a relationship held between the spectral regions of SWIR-2 and less nose-prone VNS-1. We acquired 896 pairs of green tea leaf reflectance/transmittance test data from the Greater Alishan tea country in central Taiwan. We then modeled those SWIR-2 spectra through statistical and mathematical procedures (multivariable linear regression, linear parameter multiplication, and spectral reversion) by referring to the relationships of SWIR-2 and VNS-1 of publicly available and quality-controlled international green leaf spectral reference databases. The proposed SWIR-2 reconstruction approach was verified by using another green leaf spectral database (Angers). The low differences (RMSEs) testified to the high caliber and robustness of the SWIR-2 reconstruction approach. The performance might be further improved with the availability of more comprehensive green leaf spectral databases.

## Acknowledgments

We appreciate the field and lab assistance provided by Chi-Ching Huang, Yi-Ching Hung and Zih-Yu Shen. This work was supported by the National Science and Technology Council (Taiwan) (110-2321-B-002-016, 111-2321-B-002-019, 112-2321-B-002-016), National Taiwan University (NTU) Core Consortiums Project (NTUCC-110L890901), and the Research Center for Future Earth, the Featured Areas Research Center Program, the Higher Education Sprout Project, Ministry of Education (Taiwan).

